# Sharing is not erring: Pseudo-reciprocity in collective search

**DOI:** 10.1101/258715

**Authors:** Imen Bouhlel, Charley M. Wu, Nobuyuki Hanaki, Robert L. Goldstone

**Affiliations:** Université Côte d’Azur, CNRS, GREDEG, Nice, France; Center for Adaptive Rationality, Max Planck Institute for Human Behavior, Berlin, Germany; Department of Psychological and Brain Sciences, Indiana University, Bloomington, USA

**Keywords:** Collective search, information sharing, pseudo-reciprocity

## Abstract

Information sharing in competitive environments may seem counterintuitive, yet it is widely observed in humans and other animals. For instance, the open-source software movement has led to new and valuable technologies being released publicly to facilitate broader collaboration and further innovation. What drives this behavior and under which conditions can it be beneficial for an individual? Using simulations in both static and dynamic environments, we show that sharing information can lead to individual benefits through the mechanisms of pseudo-reciprocity, whereby shared information leads to by-product benefits for an individual without the need for explicit reciprocation. Crucially, imitation with a certain level of innovation is required to avoid a tragedy of the commons, while the mechanism of a local visibility radius allows for the coordination of self-organizing collectives of agents. When these two mechanisms are present, we find robust evidence for the benefits of sharing—even when others do not reciprocate.

## Introduction

In a competitive environment, it may seem counterintuitive to share information about a new innovation (which could be copied) or the location of resources (which could be stolen). Yet unrestricted sharing of information is widely observed in both human and animal behavior. The open-source software movement is a prominent example of this behavior, where rather than privately monetizing economically viable innovations, they are shared in order to foster broader collaboration and further innovation, leading to benefits for the public good. Yet do the individuals who share also themselves benefit? What drives this behavior and how can we understand it through the mechanisms of social cooperation and collective search behavior?

In ecology, it has been observed that many species share information via mass recruitment systems when foraging for resources, whereby successful foragers send signals (e.g., sounds or pheromone trails) to help locate resources. For example, the American Cliff Swallow has a unique call used only when it finds air-borne food (Brown, Brown, & Shaffer, 1991). The call is an unrestricted social signal to other Cliff Swallows which does not only benefit kin (i.e., no kin selection mechanisms). Rather, the individual welfare of each Cliff Swallow is improved by the recruitment of peers towards the search effort, since the collective performs better at tracking prey than any lone individual. More generally, this behavior—common in different animal populations—is known as *pseudo-reciprocity* (Connor, 1986; Clutton-Brock, 2009; Torney, Berdahl, & Couzin, 2011), meaning altruistic behavior, such as sharing information, can generate “by-product” benefits that improve individual fitness without the need for explicit reciprocation. Thus, pseudo-reciprocity in Cliff Swallows is thought to be mainly driven by the sparse and dynamic distribution of resources in the environment (Brown et al., 1991), which makes individual search very difficult, although in general the mechanisms are not well understood.

### Forms of Cooperation

The study of human cooperation has revealed different levels of mechanisms that can lead to altruistic behavior (Rand & Nowak, 2013), such as direct or indirect reciprocity. *Direct reciprocity* requires repeated interactions and conditional strategies (i.e., cooperate or defect), where the evolution of cooperation depends on the likelihood of future interactions (Axelrod, 2006; Rand & Nowak, 2013). Thus, altruistic actions are explained by the expectation of reciprocity from peers in future interactions, conditional on one’s own behavior. *Indirect reciprocity* extends this expectation of future reciprocity further, through reputation-based systems, whereby cooperative behavior is viewed as a type of social signaling to third-parties (Nowak & Roch, 2007). Thus, maintaining a positive reputation can have “downstream” effects, making it more likely for third-parties to cooperate, even if there have been no previous history of interactions.

Both direct and indirect reciprocity can be understood as forms of *impure altruism* (Andreoni, 1989), whereby altruistic behavior is explained through the expectation of future reciprocation within a self-interested, utility-maximization framework, and has been applied to game theory in public good games (Andreoni, 1990; Falk & Fischbacher, 2006; Nowak & Sigmund, 2005), but less so to collective search contexts. In contrast, *pseudo-reciprocity*, can be understood as an investment in other individuals in order to acquire or enhance by-product benefits, without the need for explicit reciprocation (Leimar & Connor, 2003; Čače & Bryson, 2007). The sharing of information about the solution to a problem or the location of resources can benefit the sharer, because it can lead others to copy and improve on the original information, which can aid the original sharer as a by-product benefit.

For instance, in the case of the Cliff Swallow (Brown et al., 1991), the individual who signals for others gains *by-product benefits* through the recruitment of peers, regardless of whether or not they also signal in the future (Connor, 1995), since recruitment immediately aids in the tracking and acquisition of food^1^. Importantly, this entails that pseudo-reciprocity can operate in systems without repeated interactions or reputation systems (Torney et al., 2011), which is not the case for direct or indirect reciprocity. While both forms of impure altruism have been greatly studied in game-theory (Dufwenberg & Kirchsteiger, 2004; Fehr & Gächter, 2002; Milinski, Semmann, & Krambeck, 2002; Lotem, Fishman, & Stone, 1999; Nowak & Sigmund, 1998), there exist relatively little study of pseudo-reciprocity in human behavior (although considerably more in animal behavior Brown et al., 1991; Connor, 1986; Wilkinson, 1992).

### The Role of the Environment in Collective Search

In a collective search task, one must learn to balance the exploration of yet unknown solutions, while also exploiting existing knowledge in order to yield immediate returns. This frames the *exploration-exploitation* dilemma, where different environments can demand a different ratio of exploration to exploitation (Barkoczi, Analytis, & Wu, 2016). The relative effectiveness of different search strategies can be drastically altered by changing one of many aspects of the task environment, such as the complexity of the problem space (Mason, Jones, & Goldstone, 2008), the connectivity of the communication network (Barkoczi & Galesic, 2016; Lazer & Friedman, 2007; Mason et al., 2008; Mason & Watts, 2012), the type of social information being communicated (Wisdom, Song, & Goldstone, 2013), and the learning strategies of each individual agent (Barkoczi & Galesic, 2016). However, one aspect taken for granted in these previous collective-search paradigms, is that information is always freely given, whereas in many real world situations agents make an explicit decision to either share or withhold information. We examine how different task environments and behavioral mechanisms influence the adaptiveness of freely sharing information.

### Goals and Scope

In which environments will we find stable benefits for freely sharing information, and where will we fail to find it? We present multi-agent simulations of a competitive search task in *n*-dimensional search spaces, with both static and dynamic reward structures. Agents are given a limited search horizon to search for rewards, with different populations of agents containing different mixtures of either *sharers* or *non-sharers*. We focus on four main mixtures (All sharers, No sharers, Free-rider, and Free-giver) in order to determine for any given agent whether or not it is better to share or to withhold information from others. We use a competitive search context, whereby agents who choose the same location in the search space must split the reward between them. In the static simulation, we use a fitness landscape where the reward is a monotonically decreasing function of the distance from a randomly selected global optimum. In the dynamic simulation, the reward function is similar, but the global optimum follows a random walk, moving on each trial.

## Static Simulations

We present a mathematical framework for studying the benefits of sharing information in a competitive search context. We use multi-agent simulations to implement this framework, and present simulation results across a variety of different environmental structures.

The search task involves *k* agents searching for rewards on an *n*-dimensional environment, where each dimension *d* in 1, 2, 3, …, *n* can take any positive or negative integer value. The task takes place over *T* = 100 trials, where at each time *t* ∈ [1, *T*], all agents individually make a decision about where to search for rewards, and then whether or not to share that information with other agents (see Information Sharing). Altogether, we vary the number of agents *k* ∈ [4, 10], the number of dimensions of the search space *n* ∈ [5, 15], the innovation rate *p_c_* ∈ {0, 0.5, 1}, the visibility radius *r* ∈ {0, 1, 2}, and the mixture of sharing strategies (All sharers, No sharers, Free-rider, and Free-giver).

### Fitness function

Each search decision is represented as a vector 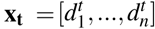, where the resulting payoff *y_t_* = *f*(**x_t_**), is given by the inverse Manhattan distance between the global optimum *M* and **x***_t_*:

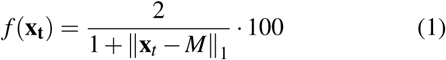

with Laplace smoothing used to prevent division by zero and rescaled by a factor of 100. The global maximum of the environment at time t is represented by *M* = [*D*_1_, …, *D_n_*], which is drawn from a uniform distribution 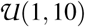.

The competitive nature of the task is represented by *splitting* earned payoffs between agents who make the same search decision on a specific trial (i.e., subtractive resources). Let **P***_t_* = [**x***_t_*_1_, …, **x***_tk_*] represent the matrix containing all *k*-agents’ search decisions at time *t*. For each agent *i*, let *C*(**x***_ti_*) be the number of rows in **P***_t_* matching **x***_ti_*,

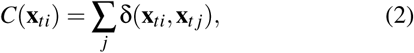

where δ(**x***_ti_*, **x***_tj_*)= 1 if **x***_ti_* = **x***_tj_*, and 0 otherwise. Thus, *C*(**x***_ti_*)= 1 when **x***_ti_* is unique, otherwise, *C*(**x***_ti_*) > 1. Thus, each agent earns reward 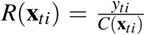, where *R*(**x***_ti_*) = *y_ti_* when no splitting of rewards occurs.

### Individual Search

At trial *t* = 1, all agents start with a random location sampled from a uniform distribution 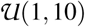. For all subsequent trials, each agent *i* has access to the history of previous observations *O_ti_*, which includes both *individually* acquired observations 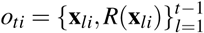 and *socially* observed data *s_ti_*, where {**x***_lj_*, *R*(**x***_lj_*)}is contained in *s_ti_* if agent *i* observed agent *j*’s (*j* ≠ *i*) information at trial *l* < *t* (see Information Sharing). Contingent on *O_ti_*, we use a local search strategy where the agent selects the search decision that had previously yielded the largest reward, 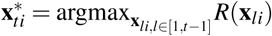, and then innovates on that solution by modifying each value in 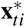 by a discrete value in {−1, 0, 1}.

We define the *Innovation rate* as the probability *p_c_* ∈ {0, 0.5, 1} with which the agent innovates on the search decision 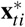. On each trial the agent innovates with probability *p_c_*, where otherwise 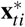 is copied verbatim. If the agent innovates, then we draw from a Binomial distribution centered on zero 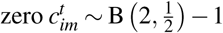 for each dimension of the search problem *m* = 1, …, *n*. Intuitively, half of the time 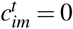, meaning there is no change along that dimension, while changes of both −1 or +1 are equally likely, each with probability 0.25. Thus, when innovation occurs, the search decision at *t* + 1 is defined as:

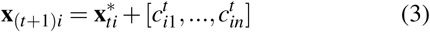

### Information Sharing

We consider four different mixtures of sharers and non-sharers in the collective. *No sharers* and *All sharers* are cases where there is a homogeneous mixture of strategies, with either all agents or no agents sharing information. In the *Free-rider* and *Free-giver* conditions, there is a single heterodox agent who acts in contradiction to the rest of the collective. The Free-rider agent is one who withholds information while all others share, and the Free-giver agent is one who shares information, even though all others withhold information. In all cases, sharing information broadcasts both the location **x***_ti_* and the earned reward *R*(**x***_ti_*) to all other agents.

We use differences in performance across these four conditions to examine the individual benefits of sharing when the behaviors of all others are fixed. By comparing the performance differences between the Free-rider agent and any given agent of the All sharer agents, we observe if there exist individual benefits of sharing when others also share. Similarly, we can compare individual performance differences between the Free-giver agent and any of the No sharer agents to observe the benefit of sharing when others don’t share. Environments where sharing is beneficial at both levels provide evidence of pseudo-reciprocity, whereby sharing confers benefits without the need for explicit reciprocation.

### Visibility Radius

In addition to explicitly shared information, we also implement a *Visibility Radius* defined as *r* ∈ {0, 1, 2} where localized clusters of agents have free access to information about each other’s current solution and rewards. For any two agents *i* ≠ *j*, agent *j* is visible to agent *i* at trial *t* if the *Chebyshev* distance between **x***_ti_* and **x***_tj_*:

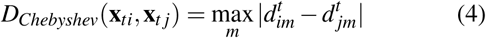

is smaller than the Visibility Radius *r*. Intuitively, Chebyshev distance is also referred as “chessboard distance”, since it is the minimum number of moves needed by a king to go from one square to another. If agent *j* is visible to agent *i* at trial *t*, then information about the current search decision and reward is automatically shared: {**x***_tj_*, *R*(**x***_tj_*)}∈ *s*_(_*_t_*_+1)_*_i_*. The visibility radius is an important coordination mechanism that allows for localized transmission of information, whereas the explicit decision to either share or withhold information operates for all distances greater than *r*. Crucially, given the high dimensionality and size of the search space, it is very unlikely for any two agents to fall within the same visibility radius without explicit information sharing.

### Results

Across each combination of environmental parameters, we conducted 10,000 simulations and computed the mean payoff over trials for each individual agent. In Figure 1 we show heat maps of the simulation results, where the color of each tile represents the net individual benefit of sharing. In Figure 1a, we compute the net benefit of sharing (when others share) as the difference between the mean payoff for a random agent from the All sharers condition and the mean payoff of a Free-rider agent. Thus, red values indicate a net benefit of sharing, while blue values indicate a net cost of sharing. In Figure 1b we run the same analysis, but compute the net benefit of sharing (when others don’t share) as the difference between the mean payoff for a Free-giver agent and a random agent from the No sharers condition. Again, red values indicate a net benefit of sharing, while values indicate a net cost. In both cases (when others share and when they do not), we find benefits for sharing information.

**Figure 1:**
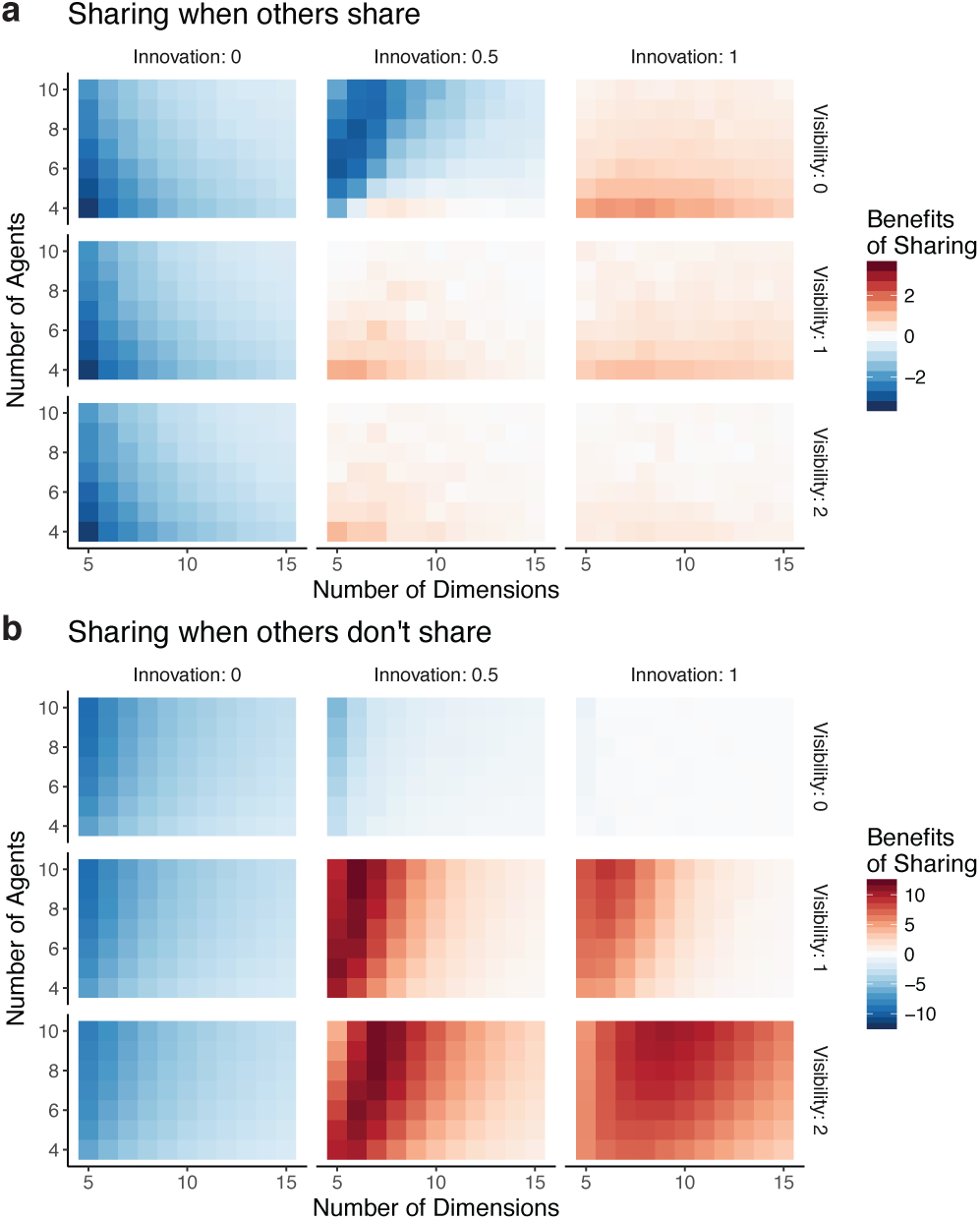
Static simulation results. We simulated 10,000 replications for each combination of environmental parameters (number of agents, number of dimensions, innovation rate and visibility radius) crossed with each of the four scenarios (all sharers, no sharers, free-rider, and free-giver). **a**) Sharing when others share. The color of each tile shows the difference in the mean agent payoffs, comparing an agent who shares when others also share (all sharers) and an agent who withholds information when others share (free-rider). Red values indicate a net benefit of sharing, white indicates no difference, and blue indicates a net disadvantage. We see that all negative side-effects of sharing disappear when both innovation and visibility are present. **b**) Sharing when others don’t share. Colors show the difference in mean agent payoffs, comparing an agent who shares when no one else shares (free-giver) and an agent who withholds information along with everyone else (no sharers). We see that both innovation and visibility support the benefits of sharing, although here we see stronger benefits, with a linear relationship between the number of agents and number of dimensions that leads to maximal benefits.

**Benefits of sharing**. Overall, we found evidence that sharing information can be a beneficial strategy, so long as peers copy with some level of innovation and in the presence of a visibility radius. Benefits were found both with and without reciprocation. Interestingly, the relative benefits of sharing when others don’t share (Free-giver vs. No sharers) were far larger than when others also share (All sharers vs. Free-rider), likely due to diminishing returns as the collective becomes more saturated with social information (see Fig 2). Thus, sharing information without explicit reciprocation leads to the largest net benefit, whereas any potential disadvantages to sharing when others share is entirely reversed by the mechanisms of innovation and the visibility radius.

**Figure 2:**
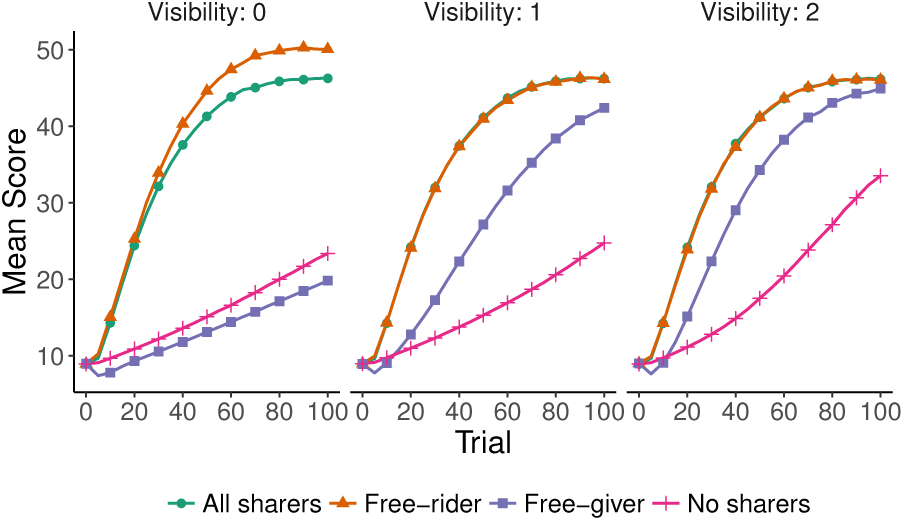
Learning curves. Mean score for 10,000 replications using *k* = 10 agents and *n* = 7 dimensions, and with innovation rate *p_c_* = 0.5. Each line is either a random agent drawn from the all sharers or no sharers condition, or specifically the heterodox agent in the free-rider and free-sharer conditions. Across different visibility radii, we see how the maladaptiveness of sharing (left panel) disappears, and transforms into an adaptive benefit. With a visibility of 1, the benefits of free-riding are reduced to zero, while the reversal for free-giving is much more dramatic and confers much larger rewards than the counter-factual case of withholding information (i.e,. no sharer).

**Number of agents and size of the environment**. As long as there are innovators (*p_c_* ≠ 0), we find that larger collectives reap larger benefits of sharing when exploring larger environments. Moreover, larger visibility increases these benefits in larger environments. Hence, the potential disadvantages of overcrowding are washed out by the advantages of social coordination through pseudo-reciprocity.

**Innovation and visibility**. Sharing information for others to imitate can be beneficial, as long as imitation also involves some level of innovation. Thus, Roger’s Paradox (Rendell & Laland, 2010), whereby social information can sometimes reduce mean fitness leading to a tragedy of the commons, only holds in our simulations when innovation is zero (Fig. 1). Additionally, Figure 2 shows the effect of the visibility radius on trial-by-trial performance, where we fix the number of agents (*k* = 10), the size of the environment (*n* = 7), and set the innovation rate to *p_c_* = 0.5. Here we see that by introducing a visibility radius, the maladaptiveness of sharing when others also share (comparing free-rider to all sharers) becomes non-existent, creating no benefit to withholding information (i.e., free-riding). Moreover, the visibility radius even more dramatically reverses the performance difference between sharing while others don’t (free-giver) and when no social learning takes place (no sharers). Whereas without a visibility radius, being a free-giver is a maladaptive strategy, we find a stark reversal when the visibility radius is introduced, where free-giving dramatically increases individual fitness, and even begins to approach the performance levels of the all-sharers condition. In our simulations, we find that both innovation and visibility are essential mechanisms that facilitate the adaptiveness of pseudo-reciprocity by creating conditions where sharing information confers by-product benefits to the individual—without the need for reciprocation.

## Dynamic Simulations

We adapted the static simulations so that the global maximum changes over time. The global maximum at time *t* is then defined by 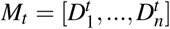. We define *Environmental change* as the probability *p_e_* ∈ {0.25, 0.5, 0.75} with which the global maximum changes at each time *t*. If the environment changes, then each dimension 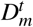 is updated using:

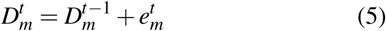

where 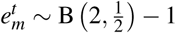, is a Binomial distribution centered on zero. Thus, environmental change operates on the same mechanism as used by agents in individual search.

In order to account for the decreasing validity of past observations in a dynamically changing environment, we introduce a *Discount rate* parameter γ ∈ {0.5, 0.6, 0.7, 0.8, 0.9} to discount the memory of previous rewards as a function of elapsed time, where smaller values of γ result in harsher discounting. Similar to temporal discounting, observations of past rewards *R_ti_* are discounted by:

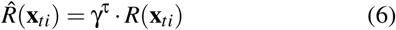

where 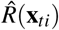 is the discounted reward observation and τ is the elapsed time between the observation and the current time.

In our dynamic simulations, we fix the number of agents and the size of the environment(*k* = 10; *n* = 7) to the combination that yielded one of the highest benefits of sharing in the static analysis, while exploring the influence of environmental change *p_e_* and the discount rate γ, along with innovation rate and visibility radius. Our goal is to find whether the sharing benefits observed in the static environment are robust to temporal dynamics in a changing environment.

### Results

Across each combination of environmental parameters and each of the four mixtures of sharers, we conducted 10,000 simulations and computed the mean payoff over trials for each individual agent, identical to the static simulations. Again, we find evidence of benefits for sharing information (Fig 3), both with and without reciprocation. While we see smaller benefits of sharing when others also share compared to the static simulations (Fig 3a), we still find that any net disadvantages of sharing are reversed into either a net benefit or a cost-free outcome when innovation and the visibility radius are present. In the case of sharing without reciprocation (Fig 3b), we find even stronger benefits of sharing than in the static simulations for particularly high innovation and visibility.

**Figure 3:**
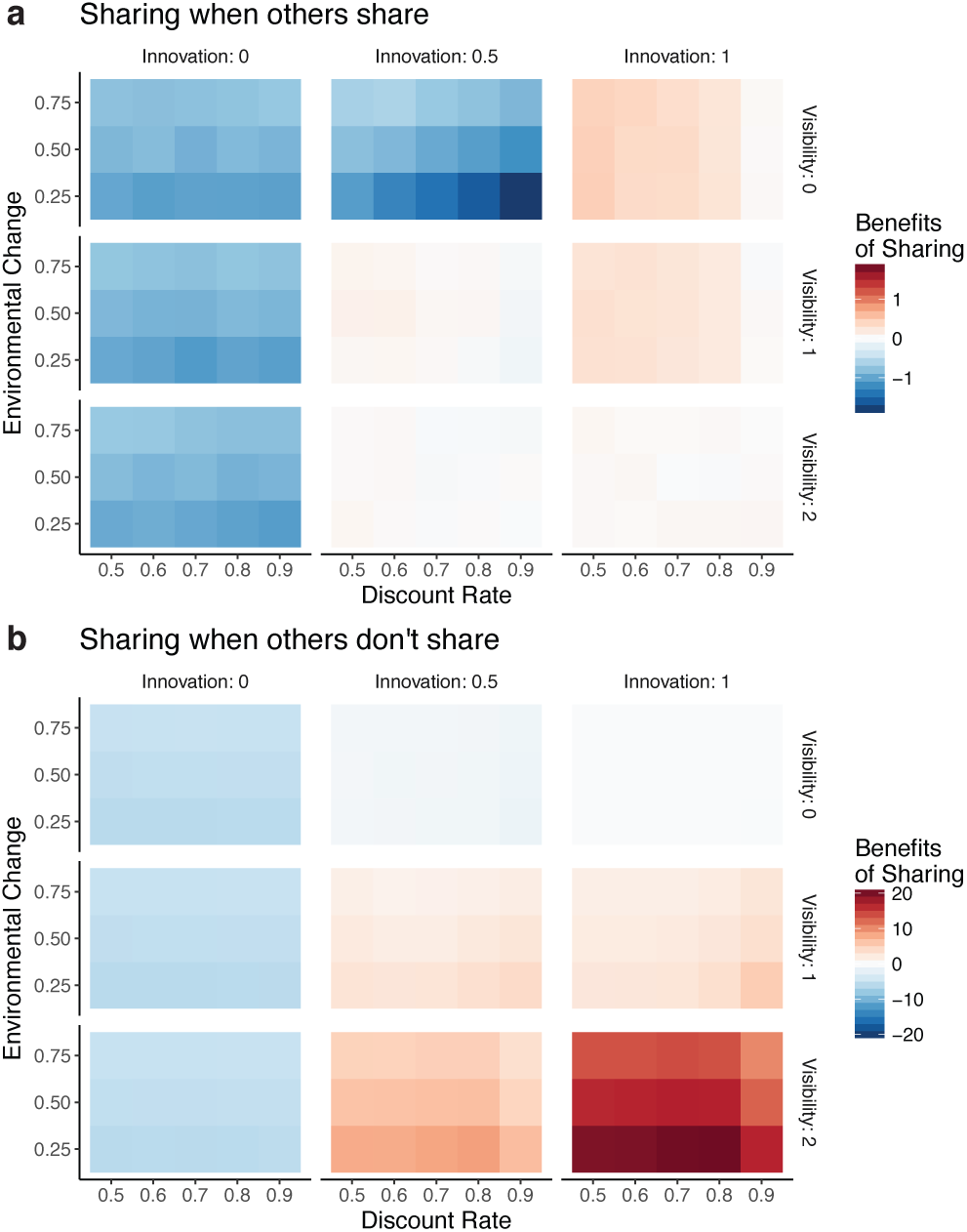
Dynamic simulation results. Here we fixed the number of agents (*k* = 10) and the number of dimensions (*n* = 7) to more freely explore differences in the dynamic environment parameters. We simulated 10,000 replications for each combination of the remaining environmental parameters (innovation rate, visibility radius, environmental change, and discount factor), crossed with each of the four scenarios (all sharers, no sharers, free-rider, and free-giver). **a**) Sharing when others share. Both innovation and visibility are robust to differences in rates of environmental change, and mediate net benefits of sharing when others also share. A strong environmental change rate also generally amplifies the net benefit of sharing. **b**) Sharing when others don’t share. The benefits of sharing are observed as soon as innovation and visibility are positively valued, and are amplified by small discount rates (resulting in faster decay rates). Moreover, with a high level of innovation (*p_c_* = 2) and visibility (*r* = 2), we see the strongest net benefits for sharing out of all simulations presented. Thus, in the dynamic simulations the local search strategy innovates on the search decision that has the largest discounted reward, 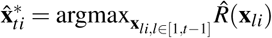.

**Environmental change and discount rate**. We observe the largest benefits of sharing in slower changing environments. Indeed, coordination of search through sharing social information is even more crucial in tracking changing rewards. However, rapidly changing environment can quickly become too difficult for any mixture of sharing strategies. We also don’t see any strong relationships between the discount rate and the rate of environmental change, suggesting that discounting past observations by itself is insufficient for adapting to a changing environment. Instead, the mechanisms of innovation and visibility played a far larger role in shaping the adaptiveness of either sharing or withholding information.

**Innovation and visibility**. We replicate the main finding from the static simulations, showing that both innovation and visibility are key ingredients for fostering pseudo-reciprocity. Each ingredient has a distinct directed effect: high innovation increases the mean performance of all agents as soon as the population contains a majority of sharers, whereas high visibility increases the individual benefits of sharing in populations where only one agent shares (Fig. 4). In dynamic environments, we find a larger benefit for sharing when others don’t share (see Fig. 4b) when we increase the visibility radius from 1 to 2 This indicates a need for more dispersed collectives of agents in changing environments, with coordination operating over larger distances. When both innovation and local visibility are present, we find that the benefits of sharing can be even larger than in the static case.

**Figure 4:**
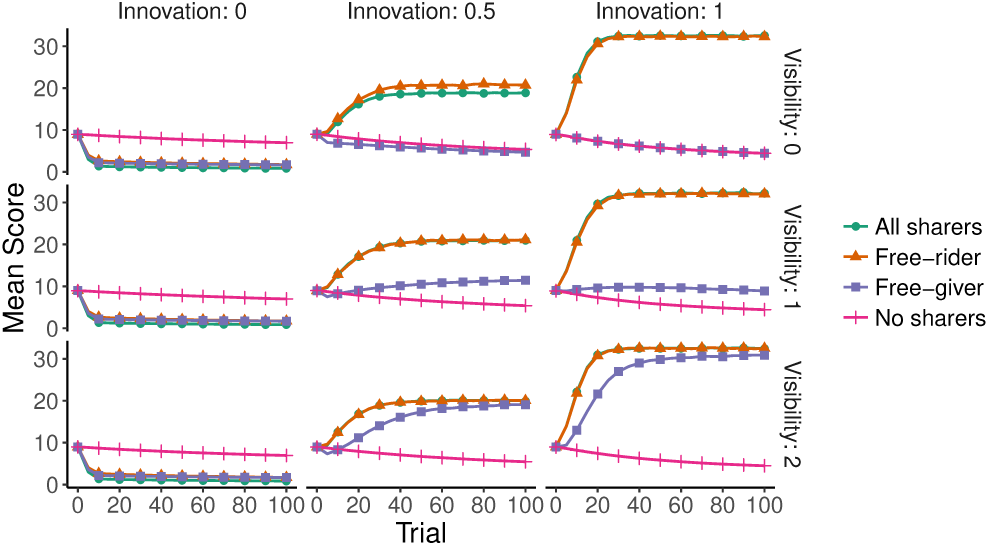
Learning curves. Mean score for 10,000 replications using *k* = 10 agents and *n* = 7 dimensions, and with environmental change *p_e_* = 0.25 and discount rate γ = 0.8. Each line is either a random agent drawn from the all sharers or no sharers condition, or specifically the heterodox agent in the free-rider and free-sharer conditions. Innovation increases the performance, and visibility increases the benefits of free giving. With positive values for both innovation and visibility radius, we find that there is never any disadvantage to sharing, either when others also share (all sharers vs. free-rider) or when others don’t (free-giver vs. no sharers).

## General Discussion

We investigated the rationality of sharing information in a competitive search context using agent-based modeling. Agents searched for rewards through *n*-dimensional environments using a local search strategy, while information about observations could either be shared with the collective or withheld. The competitive context was induced by agents being forced to equally split rewards if they chose the same location. Across both static and dynamic reward environments, we found evidence that sharing information can enhance individual performance *without the need for reciprocation*, operating on the as yet poorly-understood mechanisms of pseudo-reciprocity. Our environmental analysis of 2,700 different environments found two essential mechanisms that facilitated the benefits of sharing. Firstly, we found that sharing information for others to imitate can prove beneficial so long as imitation also involves some level of innovation. Secondly, we found that a visibility radius (allowing free information from peers within a limited distance) acted as an essential coordination device, facilitating the creation of by-product benefits for sharing. Crucially, the unrestricted sharing of information acts as a recruitment mechanism, where the visibility radius allows for small localized collectives to cooperatively find better rewards.

One limitation of our study is that we only investigate environments with a single global maximum, whereas differences in the search environments can induce different demands on the exploration-exploitation trade-off (Barkoczi et al., 2016). Indeed, highly rugged fitness landscapes may cause shared information to be a burden rather than boon, misleading individuals or collectives to explore sub-optimal local maxima of the search space. In the future, we would also look at the influence of different types of social networks governing the flow of shared information (Lazer & Friedman, 2007; Mason & Watts, 2012), along with a wider variety of individual search strategies that more closely resemble human search behavior (Wu, Schulz, Speekenbrink, Nelson, & Meder, 2018).

## Conclusion

While there have been many studies on collective search, they all typically assume free access to social information, without considering the decision to either share or withhold information. In this study, we find that even in a competitive search context, sharing information can prove beneficial, in both static and dynamic reward environments. This is one of the first studies to look at the environmental conditions that support *pseudo-reciprocity* in a competitive search context, whereby shared information can lead to by-product benefits for an individual without the need for reciprocation. This work is one of the first steps towards understanding the ecological rationality of the open source movement through the lens of a coordination system that does not require either reputation or repeated interactions.

## Footnotes

1 A crucial mechanism is that each Cliff Swallow has free, visual information about the location of nearby peers, thus allowing for localized coordination (see Visibility Radius).

